# The evolution of nitroimidazole antibiotic resistance in *Mycobacterium tuberculosis*

**DOI:** 10.1101/631127

**Authors:** Brendon M. Lee, Deepak V. Almeida, Livnat Afriat-Jurnou, Htin Lin Aung, Brian M. Forde, Kiel Hards, Sacha J. Pidot, F. Hafna Ahmed, A. Elaaf Mohamed, Matthew C. Taylor, Nicholas P. West, Timothy P. Stinear, Chris Greening, Scott A. Beatson, Gregory M. Cook, Eric L. Nuermberger, Colin J. Jackson

## Abstract

Our inability to predict whether certain mutations will confer antibiotic resistance has made it difficult to rapidly detect the emergence of resistance, identify pre-existing resistant populations and manage our use of antibiotics to effective treat patients and prevent or slow the spread of resistance. Here we investigated the potential for resistance against the new antitubercular nitroimidazole prodrugs pretomanid and delamanid to emerge in *Mycobacterium tuberculosis*, the causative agent of tuberculosis (TB). Deazaflavin-dependent nitroreductase (Ddn) is the only identified enzyme within *M. tuberculosis* that activates these prodrugs, *via* an F_420_H_2_-dependent reaction. We show that the native menaquinone-reductase activity of Ddn is important in aerobic respiration and essential for emergence from dormancy, which suggests that for resistance to spread and pose a threat to human health, the native activity of Ddn must be at least partially retained. We tested 75 unique mutations, including all known sequence polymorphisms identified among ~15,000 sequenced *M. tuberculosis* genomes. Several mutations abolished pretomanid activation *in vitro,* without causing complete loss of the native activity. We confirmed that a transmissible *M. tuberculosis* isolate from the hypervirulent Beijing family already possesses one such mutation and is resistant to pretomanid, even though it was never exposed to pretomanid. Notably, delamanid was still effective against this strain, which is consistent with structural analysis that indicates delamanid and pretomanid bind to Ddn differently. We suggest that the mutations identified in this work be monitored for informed use of delamanid and pretomanid treatment and to slow the emergence of resistance.

## Introduction

Tuberculosis (TB) is currently the leading cause of death from a single infectious agent (WHO 2016). The limitations of current treatment regimens, combined with the rapid emergence of multidrug-resistant tuberculosis (MDR-TB) strains, necessitate the development of new drugs. Three new antitubercular agents are now in advanced clinical development: bedaquiline (Andries et al. 2005) and delamanid (Matsumoto et al. 2006), which have been conditionally approved for MDR-TB treatment (Diacon et al. 2009; Gupta et al. 2015), and pretomanid (Stover et al. 2000), which is part of several promising regimens in phase III trials (http://www.newtbdrugs.org/pipeline/clinical). The nitroimidazoles, delamanid and pretomanid, are prodrugs that are reductively activated in an F_420_H_2_-dependent reaction in *Mycobacterium tuberculosis* by deazaflavin-dependent nitroreductase (Ddn) (Singh et al. 2008; Cellitti et al. 2012). An initial hydride transfer step from the F_420_H_2_ cofactor leads to their decomposition into *des*-nitro products and releases reactive nitrogen species that elicit a bactericidal mode-of-action linked to respiratory poisoning and inhibition of mycolic acid synthesis (Singh et al. 2008; Cellitti et al. 2012; Manjunatha et al. 2009).

Because pretomanid and delamanid are prodrugs that require activation, mutations that knock-out the activity of Ddn or the biosynthesis or reduction of the enzyme’s cofactor (F_420_), could confer resistance. However, the fitness cost of such knockouts may be considerable given that F_420_ has been shown to be conditionally essential to the survival of *M. tuberculosis*, being used by at least 28 different enzymes (Greening et al. 2016) and playing important roles in hypoxic survival, protection against oxidative and nitrosative damage, and evasion of the host immune system (Purwantini & Mukhopadhyay 2009; Hasan et al. 2010; Gurumurthy et al. 2013). Ddn is highly conserved across almost all species of mycobacteria (except *Mycobacterium leprae*), suggesting its physiological role is under strong evolutionary selection (Ahmed et al. 2015). It has been hypothesised that Ddn serves as an F_420_H_2_-dependent menaquinone reductase given its membrane localisation and catalytic activity with the synthetic quinone analogue menadione (Gurumurthy et al. 2013; de Souza et al. 2011), although further work is required to fully define its physiological role.

Despite the recent introduction of nitroimidazoles, cases of acquired clinical resistance, i.e. resistance that occurs during the long treatment of TB infection but is not necessarily transmissible, have already been reported (Hoffmann, Kohl, et al. 2016; Bloemberg et al. 2015). Acquired resistance to pretomanid and delamanid can occur through genetic changes that cause loss of function within the biosynthetic pathway for F_420_ production or in the F_420_-dependent glucose 6-phosphate dehydrogenase (FGD) that catalyzes F_420_ reduction to F_420_H_2_ (Choi et al. 2001; Manjunatha et al. 2006). Laboratory studies have also shown that genetic changes that abolish Ddn activity can also confer resistance (Hoffmann, Borroni, et al. 2016; Haver et al. 2015; Manjunatha et al. 2006). While such mutations could compromise treatments for already infected individuals, transmission of *M. tuberculosis* to healthy individuals after these genetic changes has never been documented. In order to spread effectively and endanger health, these resistant strains would need to retain sufficient fitness to survive all stages of the lifecycle of *M. tuberculosis,* including recovery from dormancy. Interestingly, delamanid-resistant isolates with mutations in *ddn* have been recovered from MDR-TB patients who never received delamanid or pretomanid (Fujiwara et al. 2018; Schena et al. 2016), raising important questions about the fitness costs associated with such mutations and their potential impact on transmission.

In this study, we analysed Ddn orthologs from related mycobacteria to identify natural sequence variations that make mycobacteria resistant to pretomanid, as well as analysing the genomes of ~ 15,000 *M. tuberculosis* isolates to identify the spectrum of naturally occurring non-synonymous Ddn polymorphisms. Mutations were then made to Ddn at positions identified in orthologs and through analysis of non-synonymous polymorphisms to analyse their effect on the native activity (quinone reduction) and pretomanid activation. Altogether, 75 mutants, at 47 unique positions within the 151 amino acid Ddn protein were made. This analysis identified a number of mutations that prevent pretomanid activation without full loss of the native menaquinone reductase activity. Analysis of complete and partial Ddn knock-outs in demonstrated that it is essential for resuscitation from dormancy, i.e. genetic variants that have lost their pretomanid-activation function, but retained their native activity appear to be sufficiently fit to spread and cause disease in new patients. We examined a hypervirulent strain of *M. tuberculosis* from Vietnam that contains one such mutation (despite never being exposed to pretomanid), which is resistant to pretomanid. Curiously, sensitivity to delamanid activation was not affected in this strain, nor was delamanid activation by the Ddn variant *in vitro*, which is consistent with our structural analysis that suggests it binds to the active site of Ddn in an alternative orientation.

## Results

### Ddn mutants are virulent in mice but show defective recovery from hypoxic stress *in vitro*

To understand the fitness costs of loss of Ddn activity through mutation, we investigated the ability of *M. tuberculosis* strains with mutations in Ddn to survive in stress conditions. Isogenic pretomanid-resistant *M. tuberculosis* mutants selected by pretomanid monotherapy in infected mice and shown to harbour mutations in Ddn (M1T, L49P, L64P, R112W, C149Y and from insertion of the IS6110 transposable element at D108) (Rifat et al. 2018) were used. We also analysed two mutants from a previous study in which resistant strains were identified from an *in vitro* selection experiment (S22L, W88R). We tested the ability of these eight mutants to reduce native (menadione) and drug (pretomanid and delamanid) substrates *in vitro*, finding that all mutations resulted in loss of detectable pretomanid activation, consistent with their selection in resistant strains. We observed that the M1T mutant (loss of start codon) and the mutants harbouring the IS6110 insertions did not produce any functional protein and therefore had no detectable native, nor prodrug-activating activity. In contrast, many of the point mutants, such as the L64P mutant retained a small amount of activity with menadione, suggesting that it might retain a fraction of its native function (**Table 1**).

**Table 1:**
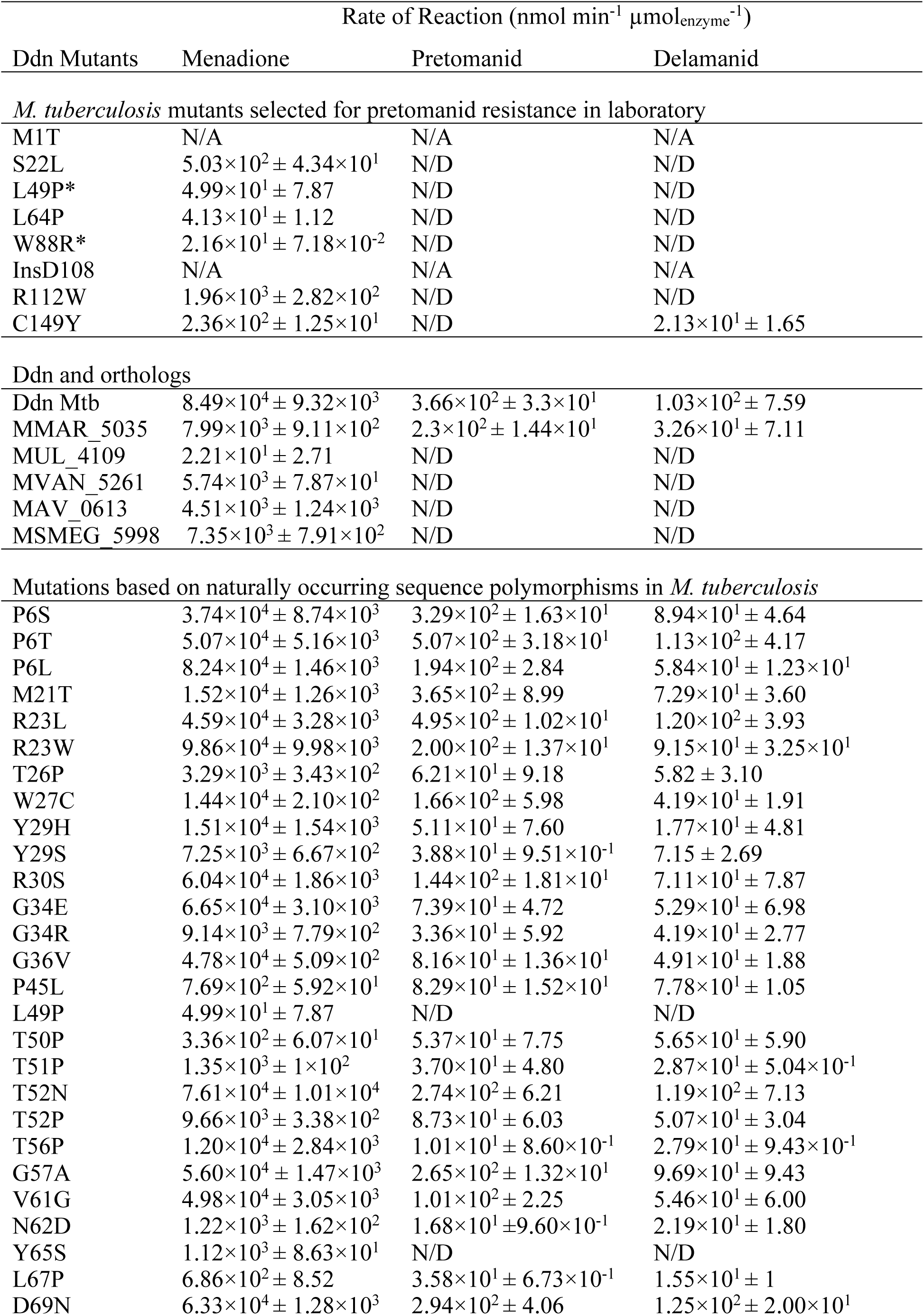
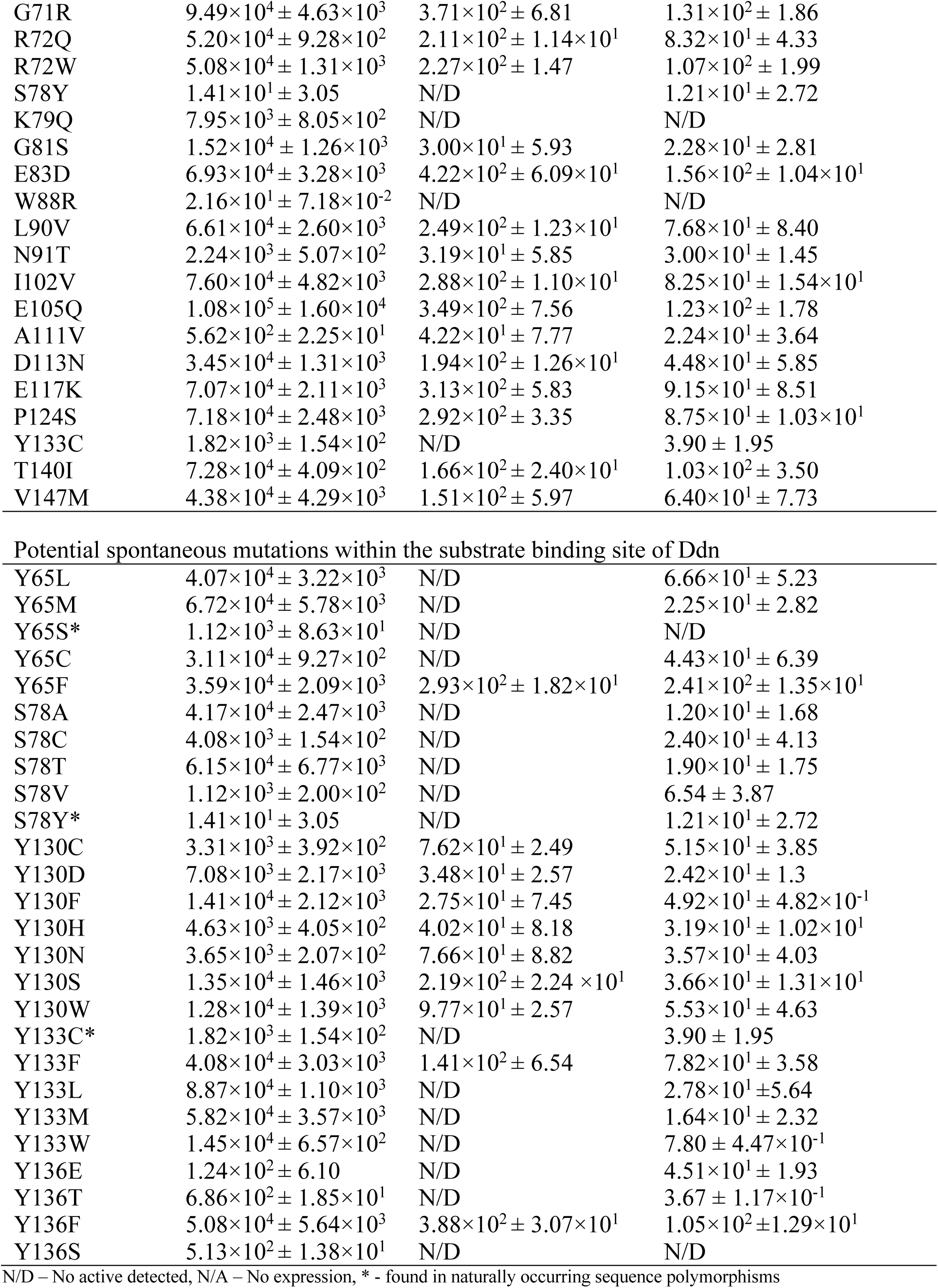
Reduction rates of F_420_H_2_-dependent menadione and nitroimidazole reductase activities of Ddn mutants.

Because many of these mutants were found in *M. tuberculosis* living in mouse lungs, the fitness cost of these mutations is clearly low enough to allow them to survive in that environment. We therefore tested whether these mutants were attenuated for multiplication and survival in mice. These mutants showed no difference in growth or survival compared to wild type after low-dose aerosol infection of mice (**Fig. 1a**), suggesting that these Ddn mutants would be transmissible. A previous study showed that *M. tuberculosis* mutants that were unable to biosynthesise F_420_ have a survival defect when recovering from hypoxia (Gurumurthy et al. 2013). We tested the Ddn mutants for their ability to survive hypoxia and resume growth upon transition to normoxia. The mutants showed no difference in survival under hypoxic conditions compared to wild type. However, they did show a significant defect in recovery from hypoxia (**Fig. 1b**) revealing the fitness cost of an inactive Ddn and providing direct evidence for the hypothesis of Gurumurthy *et al.* that Ddn has an important role in protection against oxidative stress during recovery from hypoxia (Gurumurthy et al. 2013). It is noteworthy that the L64P mutant, which retained a small amount of native activity (**Table 1**), was able to partially recover (unlike the M1T and Ins6110 mutants). Our data suggest that despite the apparent lack of fitness cost associated with loss of Ddn activity under normal and hypoxic growth conditions, dormant *M. tuberculosis* with an inactive Ddn are at a disadvantage when attempting to resuscitate, perhaps because they are more susceptible to oxidative stress or have some defect in anaerobic respiration.

**Figure 1:**
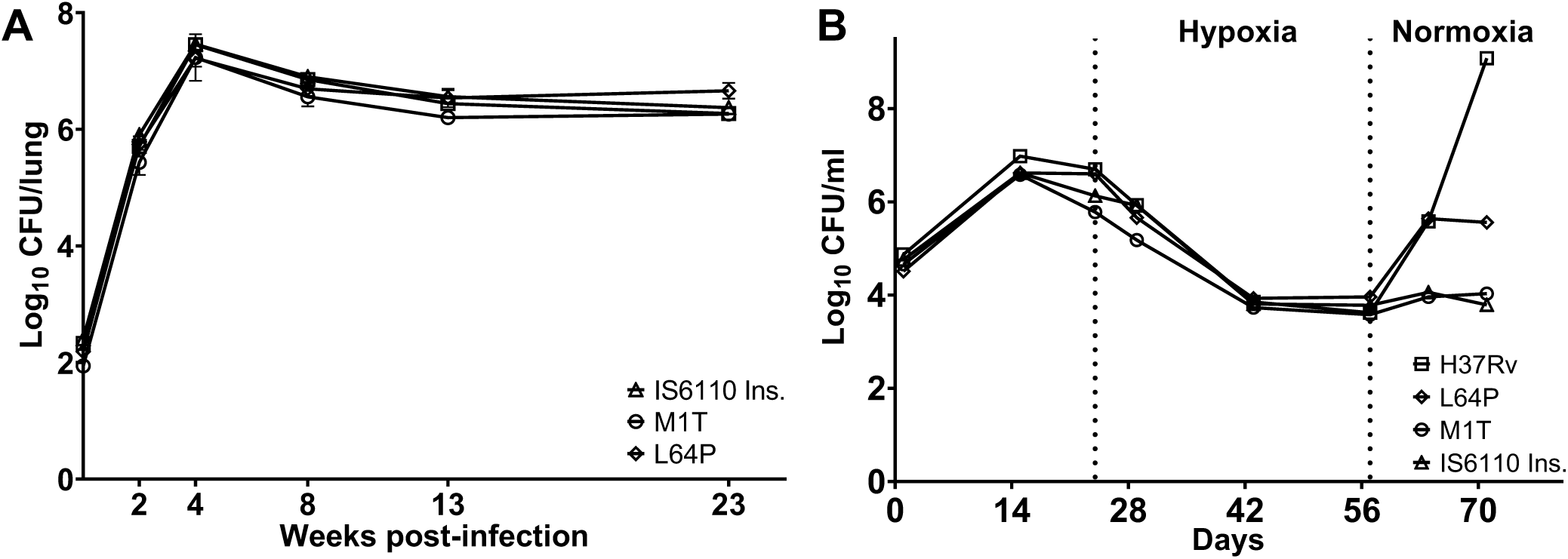
Fitness analysis of Ddn mutants. **(a)** *M. tuberculosis* Ddn mutants are not attenuated in mice. Mean lung colony-forming unit (CFU) counts of the H37Rv strain and isogenic Ddn mutants are shown following low-dose aerosol infection. Error bars represent SEM. **(b)** Isogenic *M. tuberculosis ddn* mutants have relatively normal growth and survival under progressive hypoxia but are defective in resuming growth upon return to normoxia. *M. tuberculosis* CFU counts are shown for the H37Rv strain and isogenic strains with mutations in *ddn* when grown under progressive hypoxia and resuscitated by return to normoxia. The onset of hypoxia, as defined by decolorization of the methylene blue indicator, occurred at 24 days. At 57 days, the cultures were returned to normoxic conditions.

### The physiological role of Ddn

To better understand how Ddn might contribute to recovery from dormancy we analysed its effects on aerobic respiration. A role for Ddn as a quinone reductase was previously suggested based on activity with the synthetic quinone analogue menadione (Gurumurthy et al. 2013). Here, we show that purified Ddn catalyzes the F_420_H_2_-dependent reduction of menaquinone-1, which is known to accept electrons from a variety of electron donors and transfer them to terminal oxidases or reductases in mycobacterial respiration (Cook et al. 2014). Ddn catalyzed menaquinone reduction *in vitro* with moderate efficiency (*k*_cat_/*K*_M_ = 8.6 × 10^2^ M^−1^ s^−1^) and physiologically relevant affinity (*K*_M_ = 22.4 ± 3.8 μM) **(Fig. 2A; Table 2)**; the conversion rate is likely to be higher in the native environment of the *M. tuberculosis* cell where Ddn and menaquinone are co-localised at the cell membrane (Sinha et al. 2005). Ddn orthologs encoded by *Mycobacterium smegmatis* (MSMEG_2027, MSMEG_5998) were also able to reduce menaquinone, suggesting that Ddn orthologs have similar physiological roles across the genus **(Table 2)**. Indeed, Ddn and its orthologs are highly conserved and abundant throughout mycobacteria (Ahmed et al. 2015).

**Table 2:** Kinetic parameters of native F_420_H_2_-dependent menaquinone reductase activities of Ddn and *M. smegmatis* orthologs in *in vitro* assays

| Enzyme | $k_{\text{cat}}$ (s<sup>-1</sup>) | Menaquinone | |
| --- | --- | --- | --- |
| | | $K_M$ (μM) | $k_{\text{cat}}/K_M$ (M<sup>-1</sup> s<sup>-1</sup>) |
| Ddn | $1.9 \times 10^{-2} \pm 9.7 \times 10^{-4}$ | $22.4 \pm 3.8$ | $8.6 \times 10^2$ |
| MSMEG_2027 | $5 \times 10^{-3} \pm 1.2 \times 10^{-3}$ | $80.7 \pm 34.4$ | $6.1 \times 10^1$ |
| MSMEG_5998 | $1.2 \times 10^{-2} \pm 1.5 \times 10^{-3}$ | $97.9 \pm 20.4$ | $1.2 \times 10^2$ |

**Figure 2:**
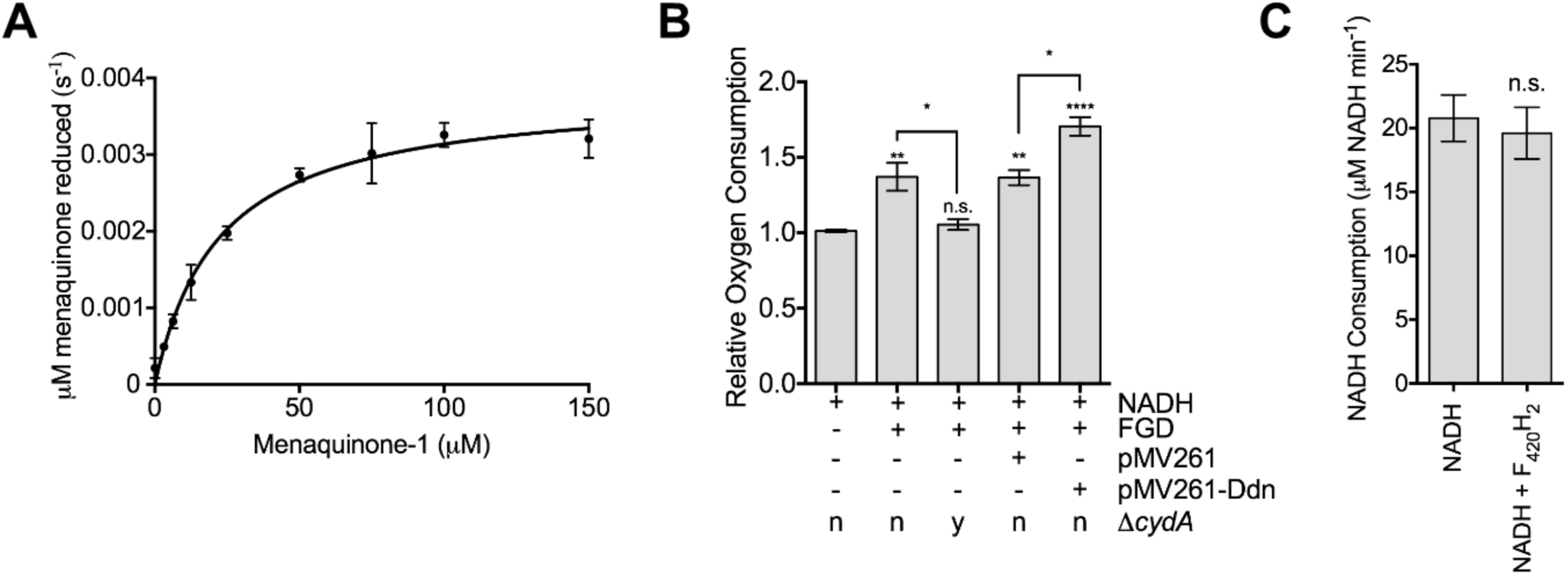
Ddn is a respiratory primary dehydrogenase that couples F_420_H_2_ oxidation to menaquinone reduction. **A.** Michaelis-Menten curve of the activity of purified Ddn with menaquinone-1 in the presence of F_420_H_2_. Rates were determined by measuring change of absorbance at 420 nm. Error bars indicate SEM (n = 3). **B.** Relative oxygen consumption of *M. smegmatis* membranes. F_420_, NADH, and glucose 6-phosphate were present in all assays, while the availability of the F_420_-dependent glucose 6-phosphate dehydrogenase (FGD) was varied. Membranes were purified from four different genetic backgrounds: wild-type, a transformant with the empty complementation vector pMV261, a transformant with pMV261 expressing the *ddn* gene, and a strain containing a chromosomal cytochrome *bd* oxidase deletion (Δ*cydA*). Error bars indicate SEM (n = 3). Stars above each column indicate a statistically significant difference compared to the first column. Other significant differences are shown by starred lines between columns. * = p ≤ 0.05, ** = p ≤ 0.01, **** = p ≤ 0.0001 n.s = p > 0.05 (one-way ANOVA followed by Tukey’s multiple comparison test). **C.** Rate of NADH oxidation in the presence and absence of F_420_H_2_. Rates were determined by following change of absorbance at 340 nm. n.s indicates p > 0.05, two tailed t-test, error bars indicate SEM (n = 3).

We then investigated whether the menaquinone reductase activity of Ddn was coupled to the mycobacterial respiratory chain by comparing the rates of respiratory oxygen consumption of the model organism *M. smegmatis* in the presence and absence of a complementation vector expressing *ddn*. We observed that the addition of glucose 6-phosphate (G6P), F_420_, and FGD, which are required to catalyze the reduction of F_420_ to F_420_H_2_ for use by F_420_H_2_-dependent enzymes such as Ddn (Oyugi et al. 2016), resulted in a 1.4-fold (P= <0.05) increase in oxygen consumption by mycobacterial membranes when the membranes are activated by NADH **(Fig 2B)**. This was dependent on the presence of cytochrome *bd* oxidase, which utilizes reduced menaquinone (i.e. menaquinol) as an electron source **(Fig. 2B)**. Thus, reduced F_420_ increases the rate of menaquinone-dependent oxygen consumption. The rate of NADH oxidation was not affected by the addition of F_420_H_2_, which implies endogenous NADH-dependent oxidases do not contribute to the increased oxygen consumption **(Fig. 2C)**. This suggest that mycobacteria can couple F_420_H_2_ oxidation to O_2_ reduction through the respiratory chain *via* cytochrome *bd* oxidase, and provides the first evidence that bacteria can use F_420_H_2_ as a respiratory electron donor. We also observed that the extent of oxygen consumption was significantly higher in membranes purified from the *ddn* expressing strain, relative to empty vector controls **(Fig. 2B)**, which is consistent with Ddn acting as a menaquinone reductase in the respiratory chain, with the remaining stimulation attributable to background activity by native Ddn orthologs of *M. smegmatis*.

### Natural sequence variation in Ddn can result in loss of pretomanid activation with retention of the native activity

Having established that Ddn is a menaquinone reductase that contributes to aerobic respiration, and that loss of this activity severely restricts resuscitation from dormancy (an essential aspect of the life cycle of *M. tuberculosis*), we then investigated the effects of natural sequence variation within Ddn orthologs from mycobacteria on the native, and drug-activating, activities. We expressed and purified Ddn orthologs from *M. tuberculosis, M. marinum, M. smegmatis, M. vanbaalenii, M. avium* and *M. ulcerans* **(Table 1)**. All Ddn orthologs catalyzed the reduction of menadione, suggesting that this is a shared physiological function that has been under consistent selective pressure. However, only Ddn from *M. tuberculosis* and *M. marinum* could activate pretomanid **(Table 1)**. This demonstrates that the native Ddn activity can exist in in the absence of nitroimidazole reductase activity, i.e. the drug-activating activity is not coupled to the native activity.

Having demonstrated that sequence polymorphisms in the active site of Ddn and its orthologs can result in loss of pretomanid reduction activity *in vitro,* we investigated whether this corresponded to differences in nitroimidazole susceptibility *in vivo. M. tuberculosis* H37Rv and *M. marinum*, which were the only two strains that encoded Ddn orthologs with *in vitro* pretomanid activation activity, were found to be susceptible to pretomanid and delamanid treatment (**Table 3**). In contrast, species in which the Ddn orthologs did not exhibit pretomanid activation activity (*M. smegmatis, M. ulcerans, M. avium*) have been shown to be naturally resistant to pretomanid (Stover et al. 2000; Ji et al. 2006; Upton et al. 2015). This analysis shows that the prodrug-activating activity of Ddn from *M. tuberculosis* H37Rv and *M. marinum* must result from sequence differences to the other Ddn orthologs tested.

**Table 3:** MICs of pretomanid and delamanid against different mycobacterial strains

| Strain | Pretomanid |  | Delamanid |  |
| --- | --- | --- | --- | --- |
|  | Reduction | MIC | Reduction | MIC |
|  | (nmol min<sup>-1</sup><br>μmol<sub>enzyme</sub><sup>-1</sup>) | (μg mL<sup>-1</sup>) | (nmol min<sup>-1</sup><br>μmol<sub>enzyme</sub><sup>-1</sup>) | (μg mL<sup>-1</sup>) |
| *M. tuberculosis* H37Rv<sup>a</sup> | 366 ± 33 | 4 | 103 ± 7.59 | 32 |
| *M. tuberculosis* N0008<sup>a,b</sup> | 0 | 256 | 12.1 ± 2.72 | 32 |
| *M. marinum* M | 230 ± 14.4 | 16 | N/A | 16 |
| *M. smegmatis* mc<sup>2</sup>155 | 0 | >50* | N/A | >50* |

<sup>a</sup>MIC was determined by reazurin assay, measuring the minimal cell death concentration <sup>b</sup>S78Y mutation
\*MICs obtained from Upton, A. et al. (Upton et al. 2015)

Genome sequences of *M. tuberculosis* were then searched to identify nonsynonymous sequence polymorphisms within the *ddn* gene. We found that around 1.5% (219/14,876) of presumptive *M. tuberculosis* genomes screened encoded a non-synonymous mutation in *ddn*, including several that have arisen independently in unrelated strains (**Table S1**). Altogether, we identified 46 non-synonymous substitutions and 2 deletions in *ddn*, distributed throughout the *M. tuberculosis* phylogeny **(Table 4)**.

**Table 4:** Distribution of Ddn alleles in the major *M. tuberculosis* lineages. Distribution of Ddn alleles in five major lineages of *M. tuberculosis.*

| Ddn allele | #Strains | Lineage(s) |
| --- | --- | --- |
| A111V | 12 | Lineage 4 (Euro American) |
| D113N | 7 | Lineage 5 (West Africa) |
| D69N | 1 | Lineage 4 (Euro American) |
| E105Q | 1 | Lineage 4 (Euro American) |
| E117K | 1 | Lineage 2 (East Asian-Beijing) |
| E83D | 4 | Lineage 4 (Euro American) |
| G34R | 11 | Lineage 4 (Euro American) |
| G34E | 2 | Lineage 4 (Euro American) |
| G36V | 1 | Lineage 1 (Indo-Oceanic) |
| G57A | 5 | Lineage 1(Indo-Oceanic),Animal Lineage |
| G71R | 1 | Lineage 4 (Euro American) |
| G81S | 9 | Lineage 2 (East Asian-Beijing) |
| I102V | 1 | Lineage 4 (Euro American) |
| K79Q | 1 | Lineage 4 (Euro American) |
| L49P | 16 | Lineage 2 (East Asian-Beijing) |
| L67P | 2 | Lineage 4 (Euro American) |
| L90V | 1 | Animal Lineage |
| M21T | 1 | Lineage 3(East African-Indian) |
| N62D | 1 | Lineage 4 (Euro American) |
| N91T | 1 | Lineage 4 (Euro American) |
| P124S | 1 | Lineage 2 (East Asian-Beijing) |
| P45L | 2 | Lineage 4 (Euro American),Lineage 1 (Indo-Oceanic) |
| P6L | 2 | Lineage 2 (East Asian-Beijing) |
| P6S | 9 | Lineage 1 (Indo-Oceanic) |
| P6T | 2 | Lineage 3(East African-Indian) |
| P86_M87del | 22 | Lineage 4 (Euro American) |
| R23L | 1 | Lineage 4 (Euro American) |
| R23W | 14 | Lineage 4 (Euro American) |
| R30S | 3 | Lineage 2 (East Asian-Beijing) |
| R72Q | 3 | Lineage 4 (Euro American) |
| R72W | 34 | Lineage 1 (Indo-Oceanic) |
| S78Y | 3 | Lineage 2 (East Asian-Beijing) |
| T140I | 1 | Lineage 4 (Euro American) |
| T26P | 2 | Lineage 4 (Euro American) |
| T50P | 2 | Lineage 2 (East Asian-Beijing) |
| T51P | 1 | Lineage 2 (East Asian-Beijing) |
| T52N | 1 | Lineage 2 (East Asian-Beijing) |
| T52P | 2 | Lineage 3 (East African-Indian),Lineage 4 (Euro American) |
| T56P | 5 | Lineage 4 (Euro American),Lineage 3 (East African-Indian) |
| TTT50PPP | 1 | East African-Indian |
| V147M | 1 | Lineage 1 (Indo-Oceanic) |
| V61G | 5 | Lineage 4 (Euro American) |
| W27C | 1 | Lineage 2 (East Asian-Beijing) |
| W88R | 2 | Lineage 4 (Euro American) |
| Y122_M129del | 5 | Lineage 4 (Euro American) |
| Y133C | 1 | Lineage 1 (Indo-Oceanic) |
| Y29del | 2 | Lineage 2 (East Asian-Beijing) |
| Y29H | 1 | Lineage 4 (Euro American) |
| Y29S | 2 | Lineage 4 (Euro American) |
| Y65S | 3 | Lineage 4 (Euro American) |
| TG56PA | 6 | Lineage 3 (East African-Indian),Lineage 4 (Euro American) |

All 46 mutants found in our genomic study were expressed, purified, and assayed with menadione, pretomanid, and delamanid **(Table 1, Fig. 3 and 4)**. Every mutant tested was able to reduce menadione, indicating that there was selective pressure to maintain this physiological function of Ddn. The majority of the mutants could also reduce/activate pretomanid and delamanid. This is not unexpected as these *M. tuberculosis* strains have not been exposed to either drug and therefore, have had no selective pressure to develop resistance. However, of the 46 mutants, several mutants did not activate pretomanid (L49P, S78Y, K79Q, W88R, and Y133C) indicating that any strain of *M. tuberculosis* with these mutations would be unable to activate the drug. Indeed, an *M. tuberculosis* strain haboring the W88R mutation has been shown to be resistant to pretomanid *in vitro* (Haver et al. 2015), and the L49P mutant was obtained from an *in vivo* resistance selection experiment (Rifat et al. 2018). It is notable that the S78Y and Y133C mutants retained the ability to activate delamanid.

**Figure 3:**
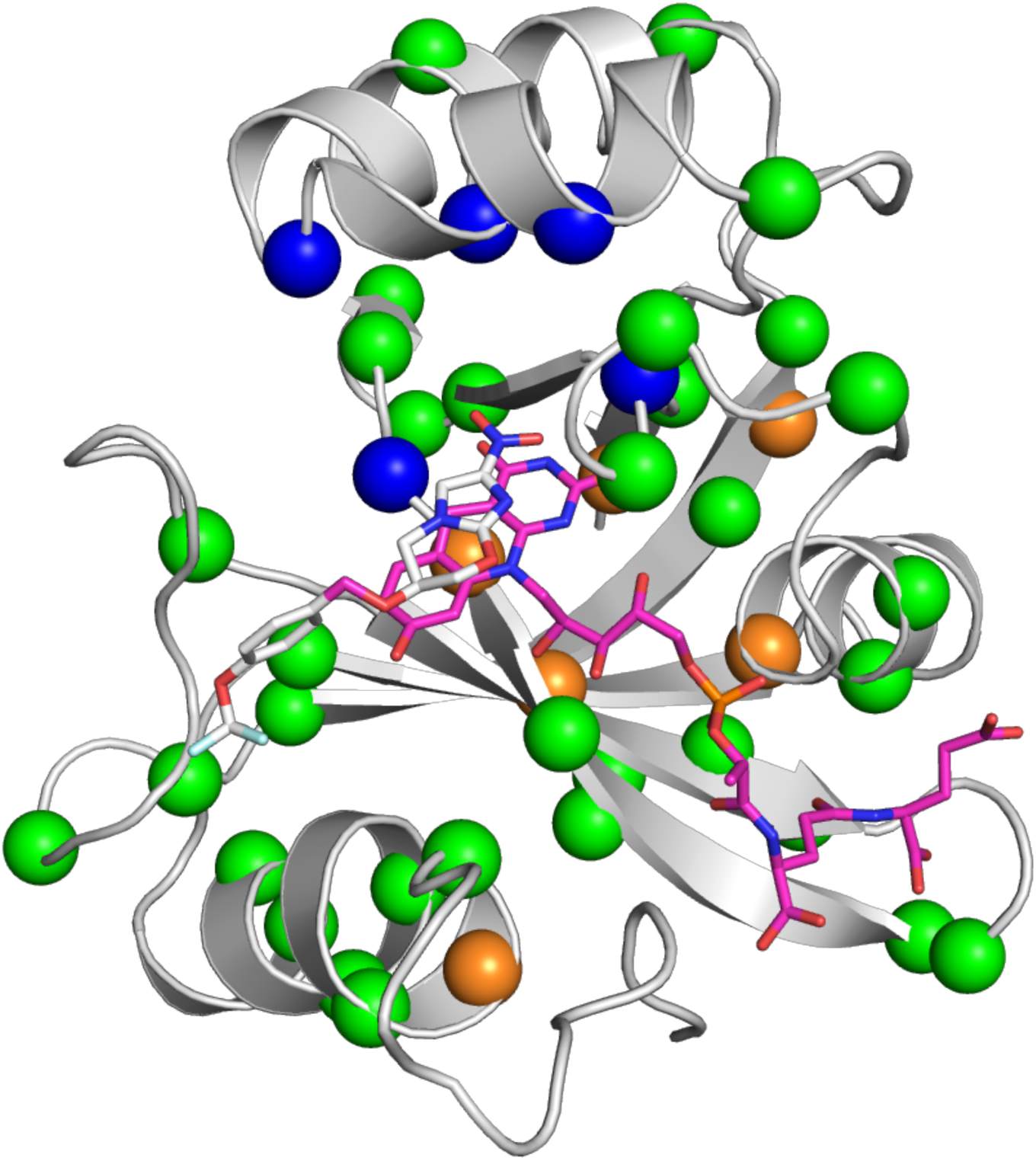
Mutations to Ddn that were made in this study. The holo-enzyme structure of Ddn with a reconstructed N-terminus and F_420_ and pretomanid bound is shown, with mutations highlighted from mutants selected for pretomanid resistance in laboratory (orange spheres), non-synonymous mutations found in sequenced *M. tuberculosis* strains (green spheres), and potential spontaneous mutations within the substrate binding site of Ddn (blue spheres).

**Figure 4:**
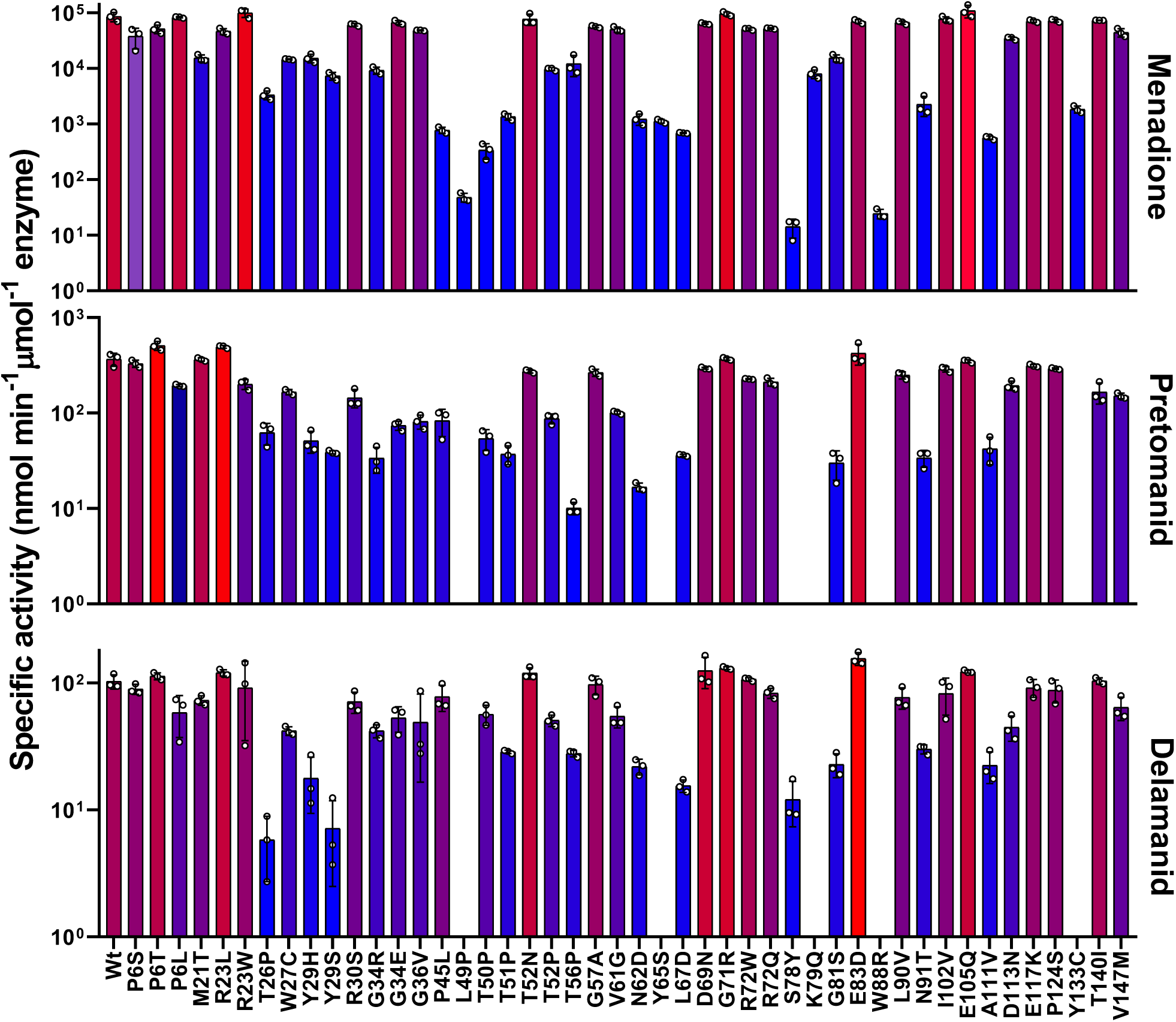
Natural sequence polymorphisms found in sequences of *M. tuberculosis* stains. The activity of Ddn mutants with menadione, pretomanid, and delamanid. Error bars show standard deviations from three independent replicates.

The S78Y polymorphism is found in the genome of N0008 (Comas et al. 2013), a clinical isolate of the hypervirulent Beijing family (Hoffmann, Kohl, et al. 2016), and in two other genomes (SRA accessions:ERR718320 and ERR751847) that are phylogenetically closely related, indicating a shared evolutionary history and suggesting that there was no substantial selective pressure to eliminate this mutation, i.e. its fitness cost must have been relatively small **(Fig. 5)**. We obtained the hypervirulent Beijing strain N0008 to investigate whether the S78Y mutation in the *ddn* gene, which results in loss of pretomanid reduction *in vitro* **(Table 4)**, corresponded to resistance to pretomanid *in vivo*. No other genetic changes previously observed to cause nitroimidazole resistance were apparent in the genome of N0008. We observed a 64-fold increase in minimum inhibitory concentration (MIC) of pretomanid against N0008 (256 μg mL^−1^) compared to H37Rv (4 μg mL^−1^). This suggests SNPs, such as L49P, S78Y and W88R, in the *ddn* gene can confer resistance to pretomanid and confirms that transmissible pretomanid-resistant populations of *M. tuberculosis* already exist.

**Figure 5:**
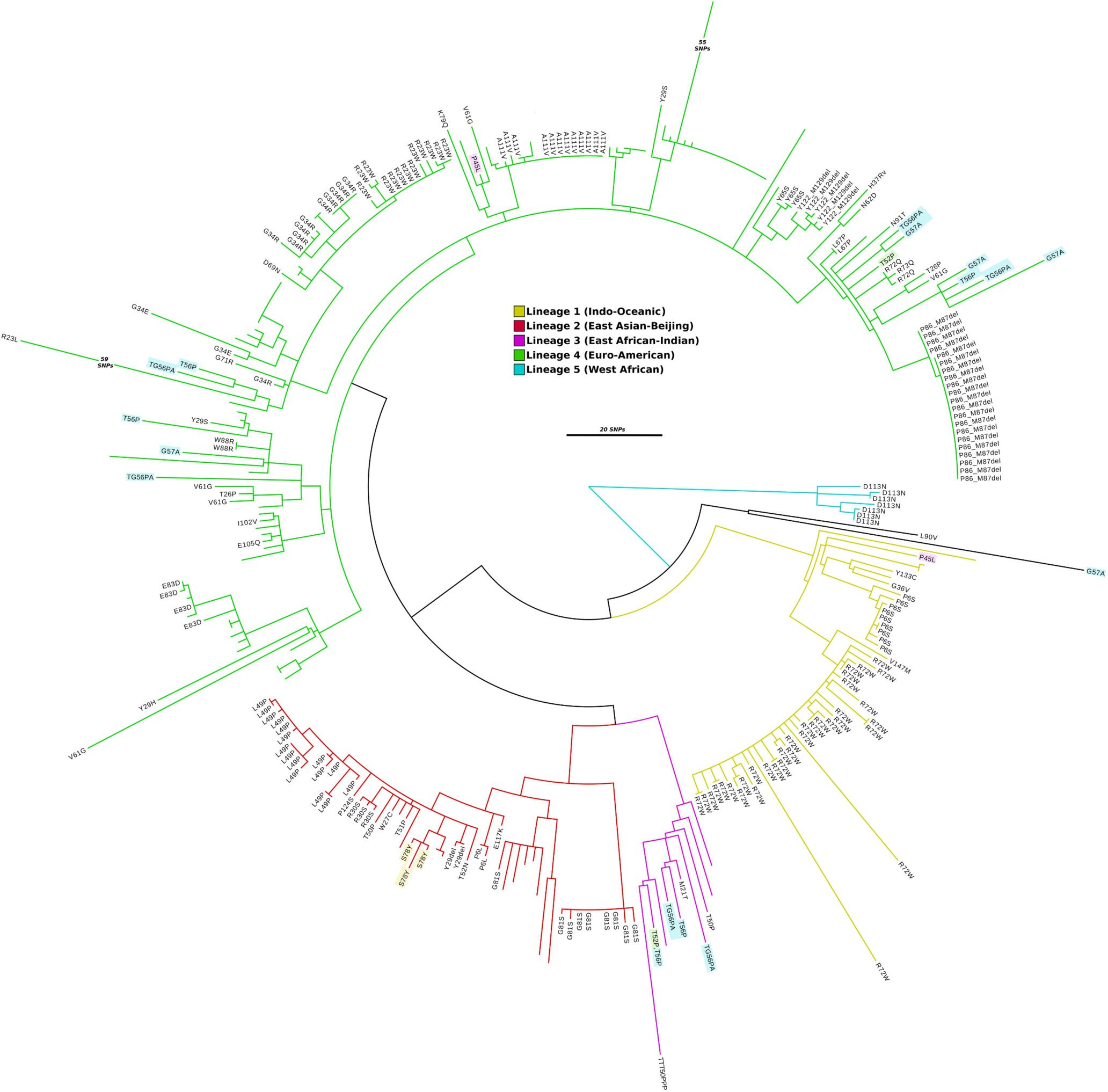
Phylogeny of *M. tuberculosis* strains with sequence polymorphisms in Ddn. Maximum-likelihood phylogenetic tree of 332 *M. tuberculosis* strains built using 2,099 non-recombinant core genome SNPs (relative to H37Rv). The tree shows all *M. tuberculosis* strains for which public genome data is available in Genbank or the short read archive (SRA) that have either synonymous and non-synonymous mutations in Ddn (relative to H37Rv). The amino acid changes of the non-synonymous mutations are indicated on the branch tips. Highlighted taxa labels indicate non-synonymous mutations that have arisen independently in unrelated strains. Blank tips represent strains with synonymous mutations in Ddn. Branches are coloured by Lineage: Lineage 1 (Indo-Oceanic), yellow; Lineage 2 (East Asian-Beijing), red; Lineage 3 (East African-Indian), purple; Lineage 4 (Euro-American), green; Lineage 5 (West African), teal. The scale bar indicates branch length in number of SNPs. Genome alignments, recombination filtering and phylogenetic reconstruction were done using Parsnp, Gubbins and RaxML, respectively. The phylogenetic tree was visualised using Figtree version 1.4.3 (https://tree.bio.ed.ac.uk/software/figtree).

### The potential for spontaneous pretomanid resistance mutations to arise in *M. tuberculosis*

Previous studies have used laboratory evolution or engineering to investigate the potential for pathogens to evolve resistance to antibiotics (Hart et al. 2016; Orencia et al. 2001). We took a similar approach here, using structure-guided mutagenesis to investigate the robustness of the nitroreductase activity of Ddn to mutations. The binding site of Ddn has been defined over several studies (Cellitti et al. 2012; Ahmed et al. 2015; Mohamed et al. 2016), identifying a number of polar amino acids within the substrate binding site that contribute to activity, particularly Y65, S78, Y130, Y133, Y136. Three of these binding site residues are fully conserved among the Ddn orthologs tested here, whereas Y65 and Y133 are more variable **(Fig. 6)**. We made, expressed and purified 26 mutants (including the Y65S, S78Y, and Y133C sequence differences observed some of the other Ddn orthologs) of these five key substrate binding residues (Y65, S78, Y130, Y133, Y136) (**Fig. 3)**, to test how spontaneous mutations at these positions affect nitroimidazole activation. Across the 26 mutants tested, all retained significant levels of the native activity, while 16 did not display detectable pretomanid activation **(Table 1, Fig 7)**. In other words, while the native activity was not greatly affected by sequence variation, the promiscuous nitroreductase activity was extremely sensitive to mutation.

**Figure 6:**
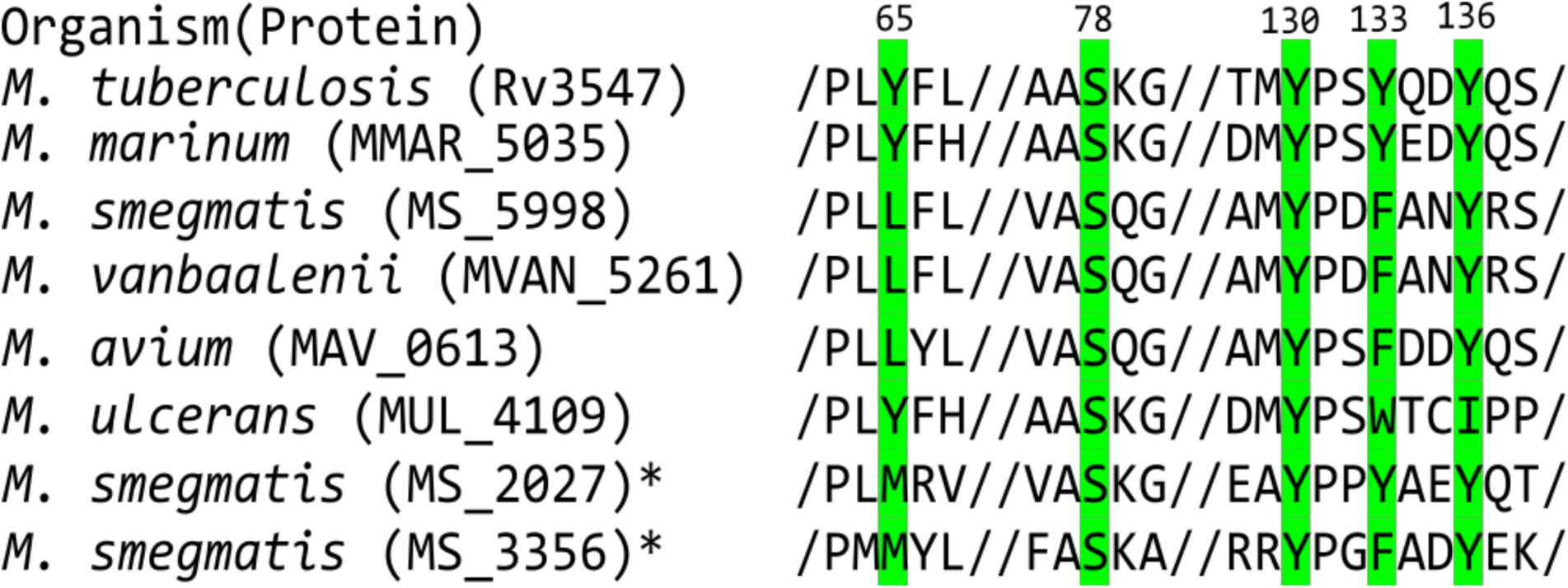
Multiple sequence alignment of Ddn and orthologs from other mycobacterial species. Highlighted residues indicate active site residues and numbers indicate their residue position in Ddn. *Indicates enzymes tested from a previous study (Ahmed et al. 2015).

**Figure 7:**
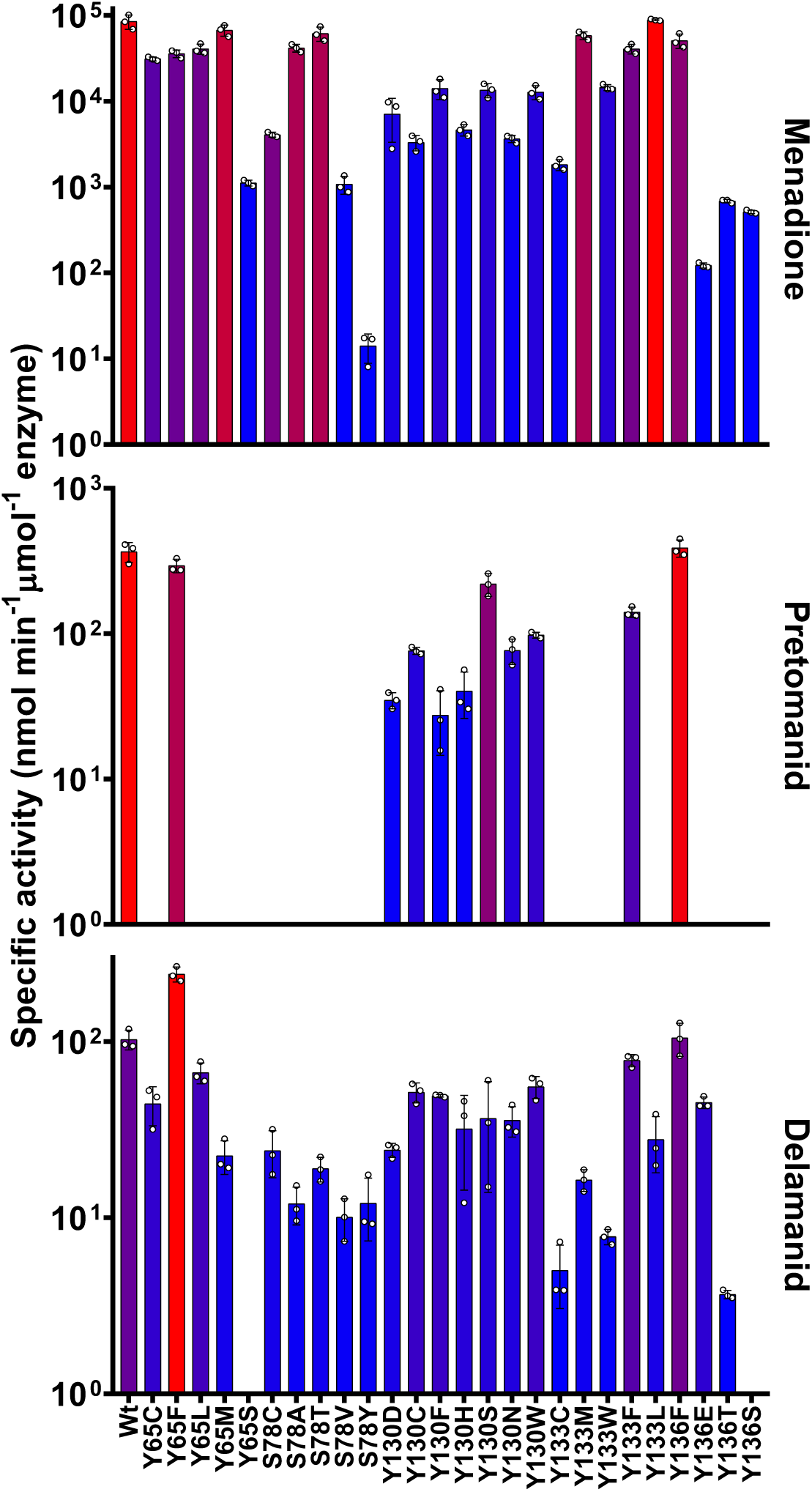
Kinetic activities of mutants with potential spontaneous mutations within the substrate binding site of Ddn. The activity of Ddn mutants with menadione, pretomanid, and delamanid. Error bars show standard deviations from three independent replicates.

To estimate the rate at which mutations could spontaneously arise at one of the positions identified in this work, we used our genomic survey of *ddn* in *M. tuberculosis* to estimate the level of sequence variation in *ddn*. This revealed that virtually every distinct lineage of *M. tuberculosis* contains SNPs in *ddn*; given the burden of TB in the world and the potential for widespread pretomanid administration, alongside the apparent sensitivity of the promiscuous pretomanid activation activity to such SNPs in contrast to the native activity, it appears that spontaneous resistance enabling mutations of *ddn* could readily occur and spread with sufficient selection pressure.

### The molecular basis of resistance

The effects of the mutations shown in **Table 1** are generally consistent with our mechanistic understanding of Ddn from mutagenesis and computational simulation (Mohamed et al. 2016; Cellitti et al. 2012). S78 is thought to interact with the nitro-moiety of pretomanid and to stabilize the transition state; none of the S78 mutants retained pretomanid nitroreductase activity, suggesting this interaction is particularly important. Y65, Y130, Y133 and Y136 are known to form a hydrophobic wall in the binding site, which can move during the catalytic cycle to shield the active site from solvent and thereby facilitate pretomanid reduction (Mohamed et al. 2016). This is in keeping with the observation that tyrosine to phenylalanine mutations at positions 65, 133 and 136 were essentially neutral, whereas substitution with other residues (Met, Leu, Cys, Trp, Thr, Ser, Glu) led to loss of pretomanid activation. Menadione reduction by Ddn is less susceptible to loss of activity through mutation, which is consistent with work showing native activities are substantially more robust to mutation than promiscuous activities (such as pretomanid activation) (Aharoni et al. 2005), as well as the observation that menadione is more chemically labile.

### Delamanid activation is less susceptible to resistance mutations

Of the 75 mutants we made and tested, 25 did not reduce pretomanid at detectable levels but only 10 lost the ability to reduce delamanid. Thus, although pretomanid and delamanid are superficially similar, they must interact with Ddn differently. These enzymatic data extend to whole cell activity, as we observed that the N0008 strain harbouring the S78Y mutation was resistant to pretomanid but remained susceptible to delamanid. For this analysis, we have focused on the wild-type protein, and the S78Y mutant to better understand the molecular basis for the differential effects of mutations on pretomanid and delamanid activation. We used molecular docking to obtain low-energy poses of these substrates in the crystal structure of Ddn. The binding of delamanid is predicted to be different than that of pretomanid (**Fig. 8C,D**), with the dual methyl and phenoxy-methyl substituents on the oxazole ring preventing delamanid binding above the deazaflavin ring of F_420_ in a ring-stacked orientation, as is seen in pretomanid, which has an oxazine ring with a single substituent in the analogous position. This results in a change in the angle at which the nitroimidazole group interacts with the cofactor that results in an increase in the distance to S78Y. Thus, there appears to be a molecular basis for the differing effects of the S78Y mutation on pretomanid and delamanid activity *in vitro* and *in vivo*.

**Figure 8.**
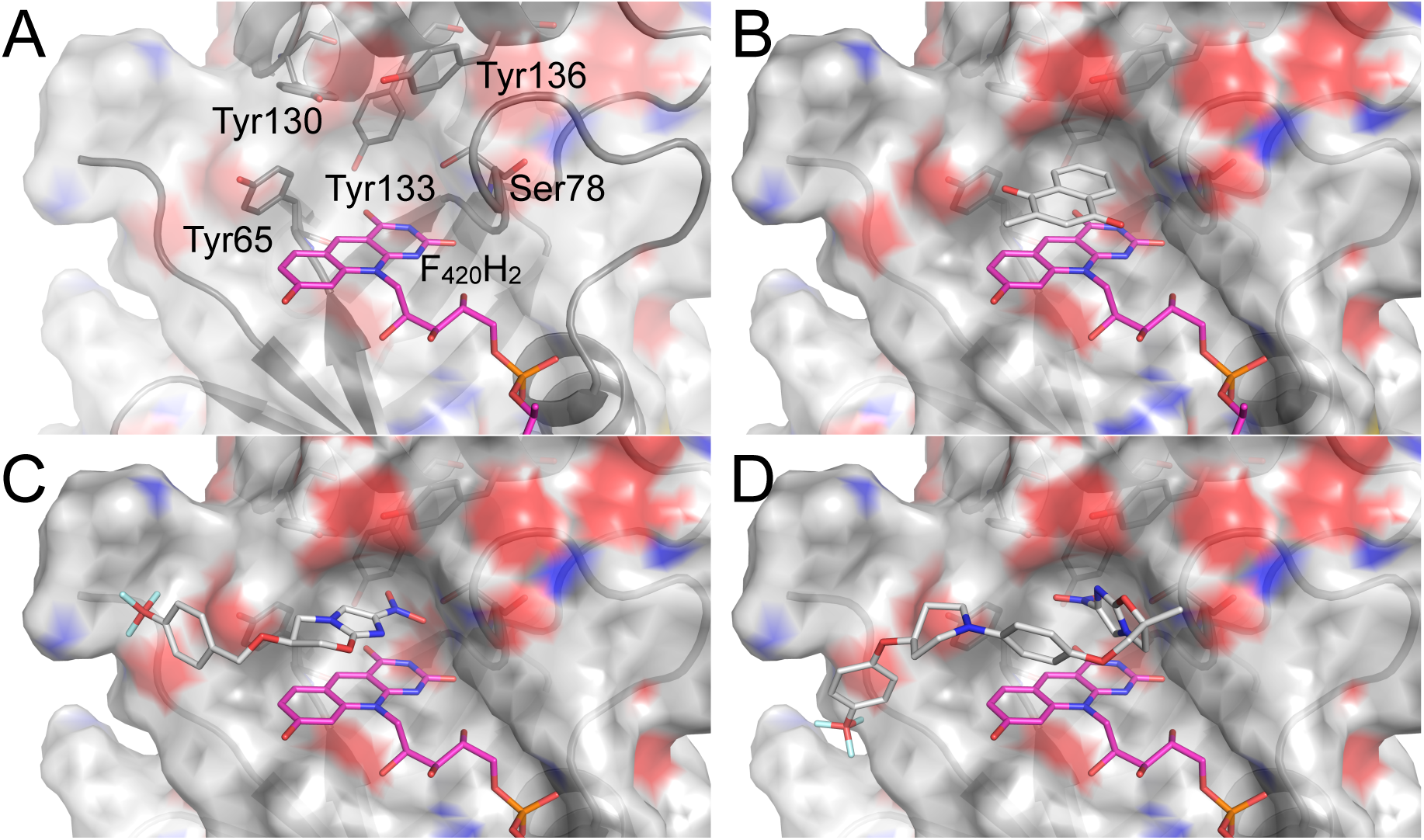
The substrate-binding pocket of Ddn (PDB: 3R5R^9^). **A.** The binding pocket of Ddn consists of F_420_/F_420_H_2_, Tyr65, Ser78, Tyr130 and Tyr136. These residues were all mutated to investigate their effects on the native (menadione) and drug-activating (pretomanid/delamanid) activities. **B.** Menadione binds above the deazaflavin ring of F_420_H_2_ in a complementary pocket formed by the aromatic rings of the tyrosine residues and the hydroxyl group of Ser78. **C.** Pretomanid docked into wild-type Ddn with F_420_H_2_ bound. The nitroimidazo-oxazine moiety can bind parallel to the deazaflavin group of F_420_, extending the nitro-group deep into the active site, towards Ser78. **D.** Delamanid docked into wild-type Ddn with F_420_H_2_ bound. The dual methyl/phenoxy-methyl substitution at the C6 position of the oxazole ring creates steric hindrance with the deazaflavin ring of F_420_, causing it to bind above F_420_ in a perpendicular orientation. This results in additional distance to Ser78.

## Discussion

### The fitness trade-off between resistance and native function

Given the conservation of Ddn-like genes throughout mycobacteria, its activity with menaquinone during aerobic respiration, and the inability of Ddn knock-outs to recover from hypoxia-induced dormancy, mutations that completely knock-out Ddn activity (including the loss of F_420_ biosynthesis through mutations to F_420_ biosynthetic genes, loss of F_420_ reductase activity through knockout of FGD, and introduction of stop codons or large genetic insertions/deletions in *ddn*) will result in substantial loss of fitness given that the ability to recover from dormancy is such an important aspect of *M. tuberculosis* pathogenesis (Diacon et al. 2009; Hards et al. 2015; Cook, Greening, et al. 2014; Lamprecht et al. 2016). Thus, for nitroimidazole resistance to spread and endanger health, the activity of Ddn must be either retained or otherwise compensated for.

The activation of the prodrugs pretomanid and delamanid by Ddn is a promiscuous activity that is not coupled to its native activity. It has long been known that promiscuous activities are more susceptible to mutation than native functions (Aharoni et al. 2005), meaning that it is possible that mutations could cause loss of prodrug activation without significant loss of native function. In this study we tested this assumption for Ddn, mutating ~ 1/3 of the amino acid positions in *M. tuberculosis* Ddn, including all known naturally occurring polymorphisms, showing that many such mutations (including several already present in clinical isolates) can cause loss of delamanid and/or pretomanid activation without loss of the native activity. Extending our enzymatic measurements to measurement of MICs revealed that related species with sequence differences in *ddn,* and clinical isolates of *M. tuberculosis* that harbor mutations within the active site, are resistant to pretomanid. These results have important implications for the clinical usage of nitroimidazoles in TB treatment in order to optimize treatment outcomes and to prevent or slow the development of resistance. For instance, pretomanid will not be effective in patients infected with naturally occurring *M. tuberculosis* variants harbouring an S78Y mutation and indiscriminant use of pretomanid against such variants (such as in regions in which the N0008 strain is endemic) could drive selective amplification and spread of pretomanid resistance. Moreover, the chances of spontaneous mutation of Ddn in currently sensitive strains is significant, given our observation of the number of single nucleotide mutations that can knock out nitroreductase activity. Given that the clinical isolate N0008 is highly transmissible, it is likely that other Ddn variants in which nitroreductase activity can be abolished without substantial loss of the native activity will also be infectious owing to the minimal loss of fitness these mutations cause with the native activity. Thus, fitness-neutral Ddn mutations are likely to be a prominent route through which clinical pretomanid and delamanid resistance spreads.

Our findings have broad implications for the continued clinical development and usage of nitroimidazole antitubercular agents. Through multiple ongoing phase II/III clinical trials, the TB Alliance and others are evaluating novel combination therapies including pretomanid. The ongoing ZeNix and TB-PRACTECAL trials are extending the study of the BPaL (bedaquiline, pretomanid, linezolid) regimen that showed highly promising results against MDR-and XDR-TB in the Nix-TB trial, while the SimpliciTB trial studies the BPaMZ regimen (bedaquiline, pretomanid, moxifloxacin, pyrazinamide) against MDR-and drug-susceptible TB. Meanwhile, comparable delamanid-containing regimens are being studied in the endTB and MDR-END trials. If delamanid and/or pretomanid are formally approved for clinical use, intensive monitoring of *ddn* polymorphisms is recommended to ensure informed regimen selection and allow interventions that will reduce spreading of transmissible resistance. Our findings indicate that delamanid binds to Ddn in a different conformation than pretomanid does, suggesting that it could be effective against some pretomanid-resistant isolates. It is also possible that combination therapy with both nitroimidazoles could help prevent the evolution and spread of resistance (since the simultaneous loss of both activities will be less likely to result from single amino acid substitution). Further studies on how delamanid and pretomanid are activated in the *M. tuberculosis* cell will inform the development of improved nitroimidazole therapies and testing of a broad range of nitroimidazole analogs against a panel of Ddn variants could help to identify compounds for which resistance is less likely to evolve.

## Materials and Methods

### Plasmid construction and point mutation

The *E. coli* codon optimised sequences for Ddn from *M. tuberculosis,* MSMEG_5998 from *M. smegmatis*, MMAR_5035 from *M. marinum,* MVAN_5261 from *M. vanbaalenii,* MAV_0613 from *M. avium,* and MUL_4109 from *M. ulcerans* were purchased as gene strings from ThermoFisher Scientific (Massachusetts, USA) and cloned into the expression vector pMAL-c2X using Gibson assembly (Gibson et al. 2009). All mutations to Ddn were made by site-directed mutagenesis using Gibson assembly (Gibson et al. 2009). Construction of MSMEG_2027 and FGD has been described previously (Lapalikar et al. 2012; Bashiri et al. 2008). For work in *M. smegmatis* the *E. coli* optimized genes were not used but instead the Ddn gene was amplified from *M. tuberculosis* DNA using the primers GTACTGCAGATGCCGAAATCTCCACCGCG, and GTAAAGCTTCTACGGTTCACAAACAACAATCGGAATG. The amplified DNA and pMV261 vector were cut using the Fast Digest enzymes PstI, and HindIII from ThermoFisher Scientific (Massachusetts, USA) and ligated together using T4 DNA ligase from New England Biolabs (Massachusetts, USA).

### Protein expression and purification

MSMEG_2027 was expressed and purified as previously described (Ahmed et al. 2015). Ddn, Ddn mutants, and Ddn orthologs were transformed into *E. coli* BL21 (DE3) cells and grown on LB agar containing 100 μg/ml ampicillin. Single colonies were picked and inoculated in LB media with 100 μg/ml ampicillin. Starter cultures were grown overnight and diluted 1/100 and grown at 37 °C until OD = 0.4. Cultures were induced with IPTG to a final concentration of 0.3 mM, and grown for 3 h at 25 °C. Cells were harvested by centrifugation at 8,500 × *g* for 20 minutes at 4 °C and resuspended in lysis buffer (20 mM Tris-Cl, pH 7.5, 200 mM NaCl) and lysed by sonication using an Omni Sonicator Ruptor 400 (2 × 6 min. at 50% power). The soluble extract was obtained by centrifugation at 13,500 × *g* for 1 h at 4 °C. The protein was purified using amylose resin (NEB) using the provided protocol. Briefly the lysate was passed over the amylose resin and washed with 12 column volumes of lysis buffer. The protein was eluted using elution buffer (same as lysis buffer but with 10 mM maltose). Samples were frozen at −80 °C in 20 mM tris pH 7.5, 200 mM NaCl, 10 mM maltose, and 10% glycerol.

FGD was expressed and purified as described by Bashiri *et al*. (Bashiri et al. 2008), with minor modifications. FGD was transformed into *E. coli* BL21 (DE3) cells and grown on LB agar containing 100 μg/ml ampicillin. Single colonies were picked and inoculated in Terrific Broth (TB) (Sambrook et al. 1989) with 100 μg/ml. ampicillin. Starter cultures were grown overnight and diluted 1/100 into auto-induction media (20 g/l tryptone, 5 g/l yeast extract, 5 g/l NaCl, 6 g/l Na_2_HPO_4_, 3 g/l KH_2_PO_4_, 6 ml/l glycerol, 2 g/l lactose, 0.5 g/l glucose, 100 mg/ml ampicillin) and grown at 30 °C for 24 h. The cells were harvested by centrifugation at 8500 g for 20 min at 4 °C and resuspended in lysis buffer (20 mM NaPO_4_ pH 8, 300 mM NaCl, 25 mM imidazole) and lysed by sonication using an Omni Sonicator Ruptor 400 (2 x 6 min at 50% power). The soluble extract was obtained by centrifugation at 13500 g for 1 h at 4 °C. The soluble fraction was filtered and loaded 5-ml HisTrap HP column (GE Healthcare) and washed with lysis buffer. The protein was eluted with elution buffer (lysis buffer with 250 mM imidazole). The purified protein was dialyzed in 50 mM Tris-Cl, pH 7, 200 mM ammonium sulphate. Samples were frozen at −80 °C in 50 mM Tris-Cl, pH 7, 200 mM ammonium sulphate, and 10% glycerol.

### Membrane purification

*M. smegmatis* mc^2^155 wild type (Snapper et al. 1990), *M. smegmatis* mc^2^155 with a kanamycin cassette disruption of the *cydA* gene (Kana et al. 2001), and *M. smegmatis* mc^2^155 transformed with empty vector pMV261 or pMV261-Ddn were all grown on LB agar supplemented with 0.05% (v/v) Tween 80. Single colonies were picked and grown in Hartmans–de Bont (HdeB) medium supplemented with 25 mM glycerol and 0.05% (v/v) Tween 80 until an OD of 0.4 to 0.8. Cultures were diluted to an OD of 0.005 in 500 ml of HdeB and grown for 72 h with agitation (200 rpm) at 37 °C. Cultures were harvested by centrifugation at, 5,000 × *g* for 15 min at 4 °C and resuspended in lysis buffer (10 mM HEPES, pH 7.5, 200 mM KCl, 5 mM MgCl_2_, 1 mM PMSF, and 1 mg DNase), homogenised on ice, and passed through a French pressure cell 3 times at 20,000 psi. Unbroken cells were removed by centrifugation at 10,000 × *g* for 10 mins at 4 °C. The membranes were isolated by ultracentrifugation at 150,000 × *g* for 45 mins at 4 °C and resuspended in working buffer (10 mM HEPES, pH 7.5, 200 mM KCl, 5 mM MgCl_2_). The protein concentrations of the membranes were measured by BCA assay (Pierce) against BSA standards.

### Enzymatic assays

F_420_ was purified from M. *smegmatis* mc^2^4517 as described by Ahmed *et al*.(Ahmed et al. 2015) F_420_ was reduced overnight with 10 μM FGD and 10 mM glucose-6-phosphate in 20 mM Tris-CL, pH 7.5 under anaerobic conditions. FGD was removed by spin filtration in a 1 mL 10 K MWCO spin filter (Millipore) and used F_420_H_2_ within 8 hours. Enzyme assays were performed according to Ahmed *et al.* (Ahmed et al. 2015; Jirapanjawat et al. 2016) in 200 mM Tris-HCl, pH 7.5, 0.1 % Triton X-100, 25 μM F_420_H_2_, and 25 μM of substrate at room temperature. Enzyme concentrations used were between 0.1 μM and 1 μM. Activity was monitored following the oxidation of F_420_H_2_ which was measured spectrophotometrically at 420 nm (ε = 41,400 M^−1^ cm^−1^) (Purwantini et al. 1992)(Purwantini et al. 1992)(Purwantini et al. 1992)(Purwantini et al. 1992)(Purwantini et al. 1992)(Purwantini et al. 1992)(Purwantini et al. 1992)(Purwantini et al. 1992). Specific enzymatic activity with delamanid was measured using fluorescence (excitation/emission: 400 nm/470 nm).

### Measurement of oxygen consumption

To determine the rate of oxygen consumption, assays were performed as described by Pecsi *et al.* (Pecsi et al. 2014b) in 10 mM HEPES, pH 7.5, 200 mM KCl, and 5mM MgCl_2_ at 37 °C with *M. smegmatis* membranes that had a protein concentration of 0.5 mg/ml, 200 μM NADH, 1 mM glucose-6-phosphate, and 25 μM F_420_ and 2.5 μM FGD where stated. The rate of oxygen consumption was measured using a model 10 Clark-type oxygen electrode (Rank Brothers Ltd., Cambridge, England) linked to a PicoLog ADC-20 data logger that was calibrated with saturated sodium dithionite as described by Pecsi et al. 2014a.

### NADH oxidation assay

The rate of NADH oxidation was determined using purified *M. smegmatis* membranes that had a protein concentration of 0.5 mg/ml in 10 mM HEPES, pH 7.5, 200 mM KCl, and 5mM MgCl_2_, 200 μM NADH, 1 mM glucose-6-phosphate, 2.5 μM FGD, and 25 μM F_420_ where stated at 37 °C. Activity was measured following the oxidation of NADH which was measured spectrophotometrically at 340 nm (6,220 M^−1^ cm^−1^).

### Computational analysis

Sequences of Ddn and orthologs were obtained from the NCBI sequence database. Alignment of the sequences were performed using MUSCLE (Edgar 2008) via the EMBL-EBI web services (Li et al. 2015). Autodock Vina (Trott & Olson 2010) was used to dock menadione, pretomanid and delamanid into Ddn (PDB ID: 3R5R) (Cellitti et al. 2012)). The protein and ligand were prepared using in Autodock tools with default settings (Morris & Huey 2009) and visualized using Pymol (DeLano 2002). Substrate structures were obtained from the ZINC database (Irwin & Shoichet 2005).

### Drug susceptibility testing

Minimum inhibitory concentration (MIC) testing of *M. marinum* was performed according to the method of Wiegand *et al*. (Wiegand et al. 2008) with some minor modifications. A culture was grown on Brown and Buckle media and colonies were scraped and diluted to an OD_600_ of 0.2, and was used as the inoculum for the MIC assay. Plates were incubated at 30 °C in a humid environment and were read after 5 days of incubation. MICs of *M. tuberculosis* isolates were determined as described previously (Heikal et al. 2016) with slight modifications. Briefly, U-bottomed 96-well microtitre plates containing Middlebrook 7H9 broth supplemented with oleic acid, albumin, dextrose and catalase were inoculated with *M. tuberculosis* H37Rv or N0008 (OD_600_ 0.02) in the presence of delamanid and pretomanid (0.015-512 μg/ml), and incubated at 37 °C for 5 days. The presence/absence of a cell pellet was checked visually. Resazurin (0.03% w/v) was then added, and plates were further incubated for 7 days. Viable cells reduce resazurin (blue) to resorufin (pink). Bedaquiline (0.015–32 μg/ml) was used as a drug susceptibility control.

### Dataset

To determine the frequency of non-synonymous sequence polymorphisms in the *ddn* gene we assembled a collection of 5,184 publicly available complete and draft *M. tuberculosis* genomes from Genbank (Benson et al. 2013) on the 7^th^ of April 2017. Additionally, using the structured query https://www.ncbi.nlm.nih.gov/sra/(Illumina[Platform]) AND “Mycobacterium tuberculosis”[orgn: txid1773]) to search the SRA (https://www.ncbi.nlm.nih.gov/sra/) <u>opened on</u> the 28^th^ of April 2017, we identified 9,692 unassembled *M. tuberculosis* genome datasets.

### Lineage-typing of *M. tuberculosis* genomes

Classification of *M. tuberculosis* phylogenetic lineages was performed using KvarQ version 0.12.3a1 and default parameters (Steiner et al. 2014). Using KvarQ scan, unassembled sequencing data were screened against a database of previously defined sequence polymorphisms known to delineate the 7 major *M. tuberculosis* phylogenetic lineages (Table S1).

### Characterisation of *ddn* sequence polymorphisms in *M. tuberculosis*

Using BLASTn (with default parameters) complete and draft *M. tuberculosis* genome assemblies from Genbank were queried with the *ddn* gene from *M. tuberculosis* H37Rv (Genbank accession:AL123456; (Cole et al. 1998)), identifying 145 *M. tuberculosis* genomes carrying alternative *ddn* alleles (relative to H37Rv). For these 145 genomes the Ddn sequence was extracted and aligned using Clustal Omega version 1.2.4 (Sievers et al. 2011) to identify non-synonymous mutation and other sequence polymorphism which could impact the function of Ddn (Table S1).

For the unassembled genomes datasets, raw sequencing reads were aligned to the reference genome *M. tuberculosis* H37Rv using Snippy version 2.9 (BWA-mem version 0.7.12 (Li 2013))(https://github.com/tseeman/snippy). Variant calling was performed using Snippy (Freebayes version 0.9.21 (Garrison & Marth 2012) with default parameters (minimum read coverage of 10x and 90% read concordance at the variant locus). 244 unassembled genome datasets were identified as having a mutation in *ddn*, only three of which were represented amongst the afore-mentioned 145 assembled genomes available in GenBank.

### Taxonomic profiling of unassembled sequencing data

Kraken (Wood & Salzberg 2014), an ultrafast sequence classification tool, was used to assign taxonomic labels to the unassembled *M. tuberculosis* genomes from the SRA. Sequence reads for each genome from the SRA were screened against a NCBI refseq database (https://www.ncbi.nlm.nih.gov/refseq) using Kraken version 0.10.5 to identify contaminants. Of the 244 *M. tuberculosis* genomes with sequence polymorphisms in *ddn*, 67 (27.5%) were found to be contaminated with DNA from unidentified organisms or from *Mycobacterium* species other than *M. tuberculosis* (Table S2). These genomes were excluded from all further analysis.

### Phylogenetic reconstruction of *M. tuberculosis* strains with sequencing polymorphisms in Ddn

To determine the phylogenetic distribution of *ddn* alleles in *M. tuberculosis* we carried out a phylogenomic analysis of *M. tuberculosis* strains with sequence polymorphism in *ddn*. Firstly, unassembled genomes from the SRA were assembled *de novo* using SPAdes version 3.9.0 (Bankevich et al. 2012). Next, all 322 complete and draft genomes (Table S3) were aligned with Parsnp version 1.2 (Treangen et al. 2014) using the genome of H37Rv as a reference to produce a core genome alignment of 848,476 bp. Core genome SNPs were identified and recombinant regions removed using Gubbins version 2.2.0 (Croucher et al. 2015). Finally, a maximum-likelihood phylogenetic tree was estimated using RAxML version 8.2.9 (Stamatakis 2014) under the GTRGAMMA nucleotide substitution model.

### Evaluation of *ddn* mutant fitness in low-dose aerosol mouse infection model

Isogenic pretomanid-resistant mutants were selected in the *M. tuberculosis* H37Rv strain background and subjected to whole genome sequencing, as previously described (Rifat et al. 2018). In the present study, female BALB/c mice were aerosol-infected with the H37Rv parent strain or one of the following isogenic *ddn* mutants, implanting approximately 2 log_10_ CFU/lung in each case: L64P, M1T, or IS6110 insertion at D108. Three mice from each group were sacrificed on the day after infection to confirm the number of CFU implanted. The multiplication and persistence of each strain was evaluated by sacrificing 3 mice/group and quantifying the CFU counts by plating serial dilutions of lung homogenate on Middlebrook selective 7H11 agar at 2, 4, 8, 13 and 23 weeks post-infection.

### Evaluation of *ddn* mutant fitness during and after hypoxic stress

The H37Rv parent and isogenic *ddn* mutants used for mouse infection were inoculated into tubes containing Dubos Tween Albumin Broth (BD Difco) supplemented with the hypoxia indicator dye, methylene blue (500 μg/ml). Tubes were sealed with rubber stoppers and incubated at 37 °C with slow magnetic stirring. The reduction and decolorization of methylene blue served as a visual indicator of oxygen levels corresponding to NRP stage 2 (Klinkenberg et al. 2008; Wayne 2001), which was attained in all cultures by day 24. Aliquots were removed for quantitative culture on 7H11 agar after 15, 24, 29, 43 and 57 days of incubation. Samples were collected by piercing the rubber stoppers with a syringe and sampling did not introduce atmospheric oxygen into the tubes, as shown by maintenance of the decolorized dye state. After day 57, each strain was returned to normoxic conditions by transferring the contents of the tubes to sterile Erlenmeyer flasks and incubated at 37 °C in a shaker for another 14 days (Sherrid et al. 2010). Aliquots were removed for quantitative culture after 7 and 14 days of normoxia.

## Footnotes

ELN reports research grants from the Global Alliance for TB Drug Development and Janssen Pharmaceuticals and collaborates on other research grants with the Global Alliance for TB Drug Development.

## Acknowledgements

We acknowledge Thomas Cuddihy (QFAB Bioinformatics) for assistance with genome data retrieval. This work was supported by by NHMRC Project Grant APP1128929 (to C. J. J., C. G., and G. M. C.). In addition, this work was supported by an ARC DECRA Fellowship DE170100310, and NHMRC New Investigator Grant APP1139832 (to C. G.). SAB is supported by an NHMRC Career Development Fellowship GNT1090456 (to SAB). DVA and ELN acknowledge the support of the U.S. National Institutes of Health (R01-AI111992).

